# What confidence and the eyes can tell about interacting with a partner

**DOI:** 10.1101/2023.02.24.529874

**Authors:** Rémi Sanchez, Anne-Catherine Tomei, Pascal Mamassian, Manuel Vidal, Andrea Desantis

## Abstract

Perceptual confidence reflects the ability to evaluate the evidence that supports perceptual decisions. It is thought to play a critical role in guiding decision-making, but only a few empirical studies have actually investigated the function of confidence. To address this issue, we designed a perceptual task in which participants provided a confidence judgment on the accuracy of their perceptual decision. Then, they viewed the response of a machine or human partner, and they were instructed to decide whether to keep or change their initial response. We observed that confidence predicted participants’ decision to keep or change their initial responses more than task difficulty and perceptual accuracy. This suggests that confidence, as a subjective evaluation of uncertainty, enables us to weigh our decisions, driving the interaction with a partner. Furthermore, confidence judgments could be predicted by pre-response pupil dynamics, suggesting that arousal changes are linked to confidence computations. This study contributes to our understanding of the function of confidence in decision-making and highlights the possibility of using pupil dynamics as a proxy of confidence.

## Introduction

Every decision we make is accompanied by a sense of confidence. For instance, when biking at night we might feel less confident about the speed and behavior of incoming traffic, and this leads us to reduce our speed. Confidence is regarded as the ability to evaluate the quality of our mental representations and decisions (Fleming & Daw, 2017). Recent theoretical accounts have suggested that confidence is critical for learning (Guggenmos et al., 2016; Meyniel, Sigman, & Mainen, 2015) as well as to enable adaptive behavior under uncertainty (Soltani & Izquierdo, 2019). For instance, it is regarded as an internal feedback signal guiding learning strategies (Bjork, Dunlosky, & Kornell, 2013) and memory “offloading” activities, such as writing down the items to buy in a shopping list (Risko & Golbert 2016). Moreover, it is also thought to be critical for human-machine interactions: poor confidence in the decisions of the machine can lead to a lack of cooperation and coordination between the human and the machine, which would hinder the effectiveness of the interaction (Zhang et al., 2020; Steyvers et al., 2022; Dafoe et al., 2021).

However, only a few empirical studies have actually investigated the role of perceptual confidence in behavior and decision-making. The difficulty in linking perceptual confidence to behavior relies on the fact that decision accuracy and confidence are strongly correlated: the higher the quality of sensory evidence, the higher the accuracy and also the confidence in one’s perceptual decisions. Hence, it remains unclear whether it is confidence or the quality of sensory evidence that influences upcoming behavior and individuals’ decision strategies (cf. Desender et al., 2018). Recent research started to tackle this issue and highlighted the contribution of perceptual confidence in information seeking behavior (Schulz et al., 2021; Desender et al., 2018) and in change of mind (Fleming et al., 2018, Rollwage et al., 2020; Pescetelli et al., 2021).

The present study aimed at investigating whether perceptual confidence, rather than first order representations of sensory evidence (i.e., linked to perceptual decisions), regulates the interaction with a partner. To address this question, participants completed a perceptual discrimination task with a partner. Participants were presented with two random dot kinematograms (RDK), presented simultaneously to the left and right of a central fixation point. They reported which RDK had a coherent motion direction closer to the vertical axis, and indicated their confidence in their perceptual decisions. Subsequently, participants viewed the perceptual decision of a (fictive) partner. They were told that the partner completed the same task and that their responses were replayed. After viewing the partner’s response, participants could keep or change their initial perceptual decision. They were explicitly instructed to use the response of their partner if they thought this could improve their performance. Importantly, to investigate whether perceptual confidence influences subsequent decisions, we needed to decouple perceptual confidence and participants’ first order representation of sensory evidence. To do this we manipulated the partner’s identity and the variability of the coherent motion. Notably, participants viewed the perceptual decision of either another human participant or a machine (more precisely., a machine-learning algorithm). They were told that the machine just like human participants was not infallible. Unknown to the participants, both partners were programmed to achieve the same level of performance. Unknown to the participants, both partners were programmed to achieve the same level of performance. We manipulated the partner’s identity, since this factor may bias confidence judgment while leaving accuracy unchanged. Indeed, recent studies showed that machines inspire overconfidence (Booth, 2017; 2020) or mistrust (Nicodeme, 2020; Lee & Rich, 2021; Seth et al., 2020) depending on the context. Secondly, we also manipulate the variability of the coherent dot motion, since it was shown that stimulus variability can distinctly impact confidence and accuracy (de Gardelle & Mamassian, 2014), i.e., high variability leads to a decrease in confidence while low variability increases confidence (cf. Desender et al. 2018). If confidence drives the interaction with a partner, we expected that participants’ decisions to change their initial judgment will more strongly be predicted by their subjective confidence rather than by their accuracy.

We were also interested in investigating the relationship between neurophysiological arousal and confidence. Arousal is an important aspect of brain function and plays a role in a wide range of behaviors, including attention, memory, and decision-making (Jing et al., 2009; Kempen et al., 2019). Recent studies have established a link between neurophysiological arousal and uncertainty (O’Connell & Kelly, 2021). For instance, changes in pupil size resulting from modulation of physiological arousal have been linked to decision uncertainty (Lempert et al., 2015; Urai et al., 2017; Balsdon et al., 2020). The ability to predict decision uncertainty from pupil dilation is important since it would allow monitoring individuals’ confidence and uncertainty continuously without explicit confidence judgments and response times. There is indeed growing interest in the field of neuroergonomics to monitor in real-time different cognitive states of operators while they interact with technology (Dehais et al., 2017, 2020; Gramann et al., 2017), and confidence appears to be a critical factor for optimal human-machine interactions (Lee & Moray, 1992; Parasuraman et al., 1993). Accordingly, this study aimed at providing evidence that pupil dilation can be used to predict individuals’ confidence judgments. However, contrary to previous studies that investigated post-decisional pupil changes linked to uncertainty, we explored pre-response pupil fluctuations that may be predictive of confidence judgments.

Finally, we also investigated the potential link between confidence and eye blink. The motivation for analyzing eye blinks was twofold: firstly, eye blinks may impact pupil dilation (Yoo et al., 2021), hence we wanted to remove this potential confound. Secondly, blinks are also thought to play a role in regulating attention and arousal: individuals tend to blink more frequently when they are engaged in tasks that require sustained attention, and eye blinks have been used as an index of an individual’s level of arousal, alertness or even fatigue (Oken et al., 2006; Maffei et al., 2018; Gavas et al., 2020).

In a nutshell, we observed that participants’ confidence predicted more reliably individuals’ decisions to change or keep their initial perceptual response than accuracy and stimulus uncertainty. Furthermore, pupil changes prior to the response predicted observers’ confidence suggesting that pupil variation can be used as an online proxy for confidence in the absence of an explicit response. Finally, eye blinks also correlated with confidence judgments. Their relation and role in decision uncertainty should be further explored.

## Materials and Methods

### Participants

Based on similar experiments investigating confidence (cf. Desender et al. 2017; Gardelle & Mamassian, 2015), the sample size was set to 14 adult participants that fully completed the experiment. In total twenty-one adults were recruited on a voluntary basis and received a financial compensation of 10€ per hour. Seven were not included in the sample size, six participants did not complete the task due to technical problems with the eyelink or difficulties with maintaining fixation during stimulus presentation, and one participant judged s/he was very confident in all her/his responses. The remaining 14 adults (9 female, range of 19 to 36 years of age) completed the experiment and were analyzed. All participants had normal or corrected to normal vision, and were naïve to the hypothesis under investigation. This study was conducted in agreement with the requirements of the Declaration of Helsinki and approved by the Ethics Committee of Université Paris Descartes. The experiment lasted approximately 2 hours and 30 minutes in total, including instructions, breaks, training, and eye-tracker calibration.

### Apparatus

Stimuli were presented on an LCD monitor (Samsung 2232RZ, 47 cm wide) with a refresh rate of 100 Hz and a resolution of 1680×1050 pixels. Stimulus presentation and data collection were performed using MATLAB with the Psychophysics Toolbox (Brainard, 1997) and the Eyelink Toolbox (Cornelissen et al., 2002). Viewing was binocular and movements of the right eye were monitored with an EyeLink 1000 Plus (SR Research, Mississauga, ON, Canada) at a sampling rate of 1000 Hz. Head movements were restrained with a chinrest located 70 cm from the screen.

### Stimuli

Stimuli were two random dot kinematograms (RDKs) presented simultaneously to the left and right of a central fixation point (a white empty circle of 0.3° visual angle diameter) at an eccentricity of 4° from fixation. RDKs consisted of 70 white dots displayed within a circular aperture of 3° visual angle diameter. Each dot had a diameter of 5 pixels (0.114° visual angle). Dots of each RDK moved coherently (5°/s speed) upward with a specific tilt angle from the vertical axis that was determined for each participant in a preliminary experiment (see below). The dot lifetime was set to 300 ms after which it was erased and then displayed in the symmetric location relative to the center of the circular aperture.

### Procedure

#### Preliminary experiment

Before starting the main experiment, we assessed participants’ tilt discrimination thresholds. Each trial started with the presentation of a fixation point. Two RDKs were displayed on either side of fixation, 500 ms after fixation onset, and for a duration of 1200 ms. The dots on the left RDK moved upward with a slight tilt toward the left, while the dots on the right RDK moved upward with a slight tilt toward the right. While keeping their gaze on the fixation, participants had to report in which RDK (left or right) the global motion moved closer to the vertical axis, by pressing the left or right arrow key of a keyboard with their right hand. Participants were told that accuracy was more important than rapidity when reporting their perceptual decisions. If fixation was broken or a blink occurred while the stimulus was displayed, an error message appeared on the screen and the trial was interrupted. Participants responded only after the RDKs disappeared and four unfilled black squares (with a side length of 0.3° visual angle) appeared around fixation (eccentricity of 1.5° visual angle) in a cross-like shape (i.e., one square above, one below, one to the left and one to the right of fixation, see Figure 1). Participants’ response was shown with the corresponding square filled with a light gray color.

**Figure 1:**
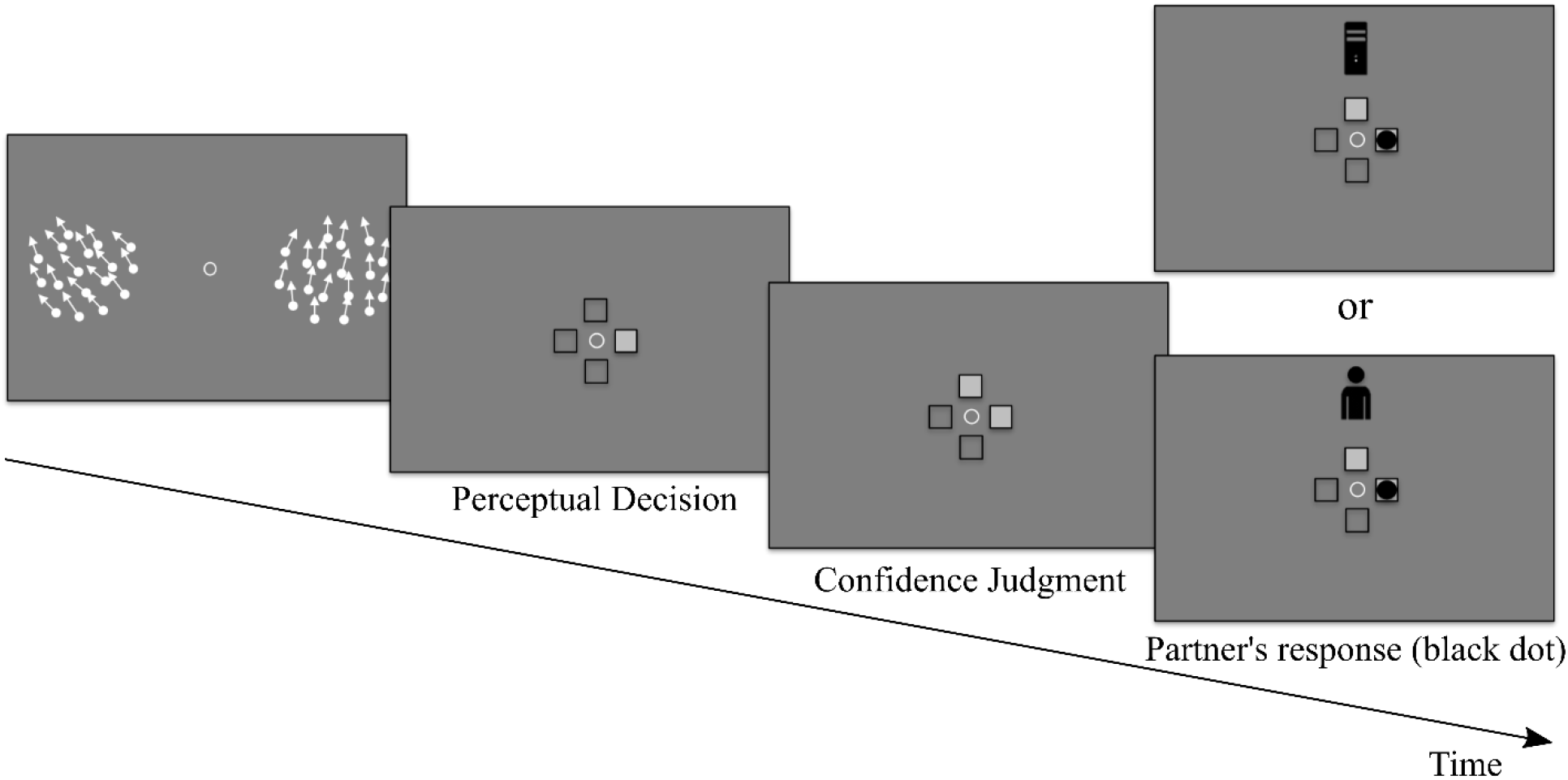
Schematic illustration of a trial. The two RDKs appeared 500 ms after a display screen that contained only a central fixation point. The dot motion was presented for 1200 ms. Thereafter, the RDKs disappeared and four squares were presented indicating the participants to report which RDK contained the dot motion that was closer to the vertical axis by pressing the left or right arrow. Participants could then report their confidence before viewing the response given by their partner. After the presentation of the partner’s response, they could choose whether to keep or change their initial perceptual judgments.

The calibration to the participants’ discrimination threshold consisted in controlling the tilt angle of the dot motion direction. In each trial we varied the tilt angle of one RDK while we kept the tilt of the other RDK to 40° (reference). The change of tilt of each RDK was controlled by two interleaved staircases one starting with a difference of 15° and the other with a difference of 30° from the reference value (i.e., 40°). If the participant’s response was correct (incorrect) the tilt difference between left and right RDK decreased (increased) in the subsequent trial. The size of this increment/decrement was controlled by an accelerated stochastic approximation algorithm (Kesten, 1958) set to converge at the tilt difference that supported 75% accuracy. Each staircase stopped once the convergence level was reached. The convergence values reached by the two staircases controlling the left (and right) RDK were averaged and used as the participant’s tilt discrimination threshold for the left (and right) RDK. Importantly, in the preliminary experiment we also manipulated the variability (standard deviation of the Gaussian distribution) of the dot motion directions. Dots could move coherently in the specified direction with a small (i.e., 3°) or a large variability (i.e., 15°). Separate tilt discrimination thresholds were obtained for these two variability levels. In sum, the preliminary experiment allowed estimating, for each participant, 4 tilt discrimination thresholds: 2 sides (left and right) × 2 variabilities (high and low).

#### Main experiment

Participants completed the same tilt discrimination task described above. The tilt difference between left and right RDKs could take one of three possible values: 70%, 100% or 140% of the tilt discrimination threshold determined in the preliminary experiment. Hence, participants completed the task under three difficulty levels: hard (70% of threshold), intermediate (100% of threshold), and easy (140% of threshold). Finally, as for the preliminary experiment, we manipulated the variability of coherent motion, i.e., dots could move coherently with a variability of 3° or 15°. We used different variability values and difficulty levels to prompt individuals to vary their confidence judgments.

After reporting the RDK with the dot motion direction closer to the vertical axis, participants indicated their confidence on a 2-point scale, that is whether they were confident or not about their response, by pressing the up or down arrows of the keyboard, for “I am sure, it was correct” and “I am not sure”, respectively. Similarly, to the perceptual decisions, participants’ confidence judgment was shown by filling the corresponding upper square or lower square (see Figure 1). Confidence judgments were followed by the presentation of the perceptual response of a machine or human partner. To remind the partner who they were interacting with, a black icon of a computer or human was displayed on top of the fixation. The partner’s judgment was presented with a black circle (0.3° visual angle diameter) appearing within the square corresponding to the perceptual judgment of the partner (left/right). After visualizing the partner’s judgments, participants decided whether to keep or change their initial perceptual response. They were instructed to use the response of their partner if they thought this could improve their discrimination accuracy. They were told that both the human and machine partner viewed the same stimuli and their answers were stored and replayed. They were also told that the machine partner was a machine learning algorithm and just like a human participant it was not infallible. In reality partners’ responses had been programmed by the experimenter and both partners were set to achieve the same performance accuracy. In particular, they were programmed to have 80, 90 and 100% accuracy for the hard, intermediate and easy difficulty level, respectively. Hence, partners’ accuracy was programmed to be higher than participants’ theoretical accuracy (75% threshold in the preliminary experiment). Human and machine partner trials were divided into 4 blocks and were presented alternately with the order of presentation counterbalanced across participants. Each block consisted of 144 trials, with Variability and Difficulty trials being randomized and equiprobable within each block. In total, participants completed 576 trials (48 trials × 3 difficulties × 2 variabilities × 2 partners).

Participants were instructed to maintain fixation and to not blink both during RDKs presentation and before their perceptual response. Fixation was considered broken when the gaze moved more than 1.25° away from the fixation center. If a fixation break or a blink occurred, the trial was interrupted, an error message was displayed (“please fixate”) and the same trial repeated at the end of the block.

### Behavioral analyses

Linear mixed models were fitted using the ‘lme4’ package (Bates et al., 2015) and the lmerTest package (Kuznetsova et al., 2017) in R (R Core Team, 2018). Factors with two levels were coded using sum contrasts (Schad et al., 2020). The models were adjusted to accommodate convergence issues. Specifically, the parsimony principle guided the selection random effects (Bates et al., 2018). Indeed, for some analyses, the inclusion of the maximal random structure led to convergence failures. In these cases, we performed a Principal Component Analysis (PCA) to isolate the random effects that contributed the least to model fitting and we removed them from the final model. Significance of the fixed effects was determined using type-II Wald tests with the ‘car’ package in R (Fox & Weisberg, 2011). Post-hoc comparisons were performed with the package ‘emmeans’ (Searle et al., 1980). When required, a False Discovery Rate (FDR; Benjamini & Hochberg, 1995) correction for multiple comparisons was applied. We used an alpha level of 0.05 for all statistical tests.

### Pupil analyses

Raw pupil data were firstly filtered with a high-pass non causal Finite Impulse Response (FIR) filter of 1/12 Hz^1^ in order to remove slow oscillations in the pupil response not related to the task. We then extracted two types of segments: 1) from -100 to +1200 ms time-locked to stimulus onset, and 2) from -1400 to +400 ms time-locked to perceptual response onset. Individual segments were corrected with a 100 ms baseline. Specifically, we subtracted the average pupil size observed from -100 ms to 0 ms prior to stimulus onset for each trial from each time point of the corresponding trials. Blinks were linearly interpolated. The correct interpolation of blinks was verified through visual inspection of each segment. Visually inspecting the segment allowed us also to remove trials containing artifacts (e.g. trials containing blinks that were not correctly labeled and interpolated, epochs containing artifactual activity such as sudden spikes of pupil changes, epochs exhibiting pupil changes that exceeded a threshold of 700 units^2^ from the baseline). This led to the removal of an average (across participants) of 2 trials in the stimulus-locked segments (i.e., 0.35% of the trials) and to 8.64 trials in the response-locked epochs (i.e., 1.5% of all trials). Pupil segments were then down-sampled by averaging the pupil size of consecutive 100 ms time windows. From the clean segments, we computed pupil dilation velocity (*v* = Δ*s*/Δ*t*; where *s* is pupil size and *t* is time) since recent studies suggest that temporal derivative of pupil size reflect more closely the dynamics of arousal fluctuation (Crombie et al., 2021; Okun et al., 2019; Reimer et al., 2014, 2016).

Pupil data were analyzed in two steps. Firstly, we performed a cluster-based permutation test comparing high confidence and low confidence trials. The aim of this first step was to isolate a temporal cluster where the pupil could dissociate low and high confidence trials. After isolating these temporal windows, pupil data points of the time window were averaged and analyzed with linear mixed-effect models to evaluate their relationship to predictors of interest.

### Classification analyses

In a series of analyses, we used Linear Discriminant Analyses (LDA; cf. Carlson et al., 2003) to evaluate to what extent pupil velocity measured during stimulus presentation (hence prior to participants’ perceptual responses) could predict individuals’ judgment of confidence. We then compared the prediction accuracy of the pupil classifier to the accuracy observed with a classifier predicting confidence judgments from individuals’ response times (i.e., confidence strongly correlates with perceptual decision latencies, notably the slower a perceptual decision the lower the confidence, cf. Mamassian, 2016). This comparison would help us evaluate the quality of pupil-based classifier predictions with respect to a well-established proxy of confidence.

LDA classifiers were trained and tested for each time-point of the pupil velocity data to dissociate high and low confidence trials for each participant. Given that the stimulus-locked segment was not contaminated by blinks, we used this time window for the classification. The classification procedure implemented a Monte Carlo cross-validation method (Dubitzky et al., 2007). Notably, each classifier was trained on 90% of the available dataset and tested on each of the remaining trials. This procedure was repeated 1000 times. Each time a random 90% of trials was used as training set and the rest as test set. To avoid classification biases at each repetition of the procedure we matched the number of high and low confidence trials to classify. Classification accuracy was estimated by calculating for each time-point the proportion of trials that the classifier correctly identified as high or low confidence trials. The mean classification performance of the 1000 shuffling within each participant and time-point was taken as the classifier accuracy for that specific participant and time-point. For statistical analyses chance level (a probability of 0.5) was subtracted from classification accuracy. Statistical significance (α = 0.05) was then calculated using a cluster-based permutation test performed on the resulting values. The same classification approach was used to classify high and low confidence trials from reaction times, except that for this variable the statistical significance was assessed with a one-sample two-tailed Wilcoxon signed rank test on the resulting values.

## Results

### Confidence

Linear mixed models assessed the influence of the Difficulty, Variability, Partner and Accuracy factors, and how these interact with confidence. Difficulty, Variability, Partner and Accuracy were also included as uncorrelated by-participants random slopes and intercepts. A main effect of Difficulty was found (χ^2^(2) = 19.61, p < 0.001; see the unsorted line of Figure 2a), with confidence decreasing with the increase of task difficulty (easy trials: M = 0.67, SD = 0.14; intermediate trials: M = 0.61, SD = 0.17; hard trials: M = 0.55, SD = 0.19). A main effect of Accuracy was also found (χ^2^(1) = 134.24, p < 0.001), with lower confidence for incorrect (M = 0.42, SD = 0.18) compared to correct judgments (M = 0.67, SD = 0.16), showing that participants could evaluate the correctness of their responses. The interaction between Accuracy and Difficulty was significant (χ^2^(2) = 34.47, p < 0.001; Figure 2a). Simple main effect analyses showed that confidence increased with the decrease of task difficulty only in correct response trials (p ≤ 0.010). There was no main effect of Variability, but the interaction between Variability and Difficulty was significant (χ^2^(2) = 13.33, p = 0.001; Figure 2c). Simple main effect analyses showed that confidence decreased in low Variability trials compared to high Variability trials only for the hard difficulty level (p = 0.014).

**Figure 2.**
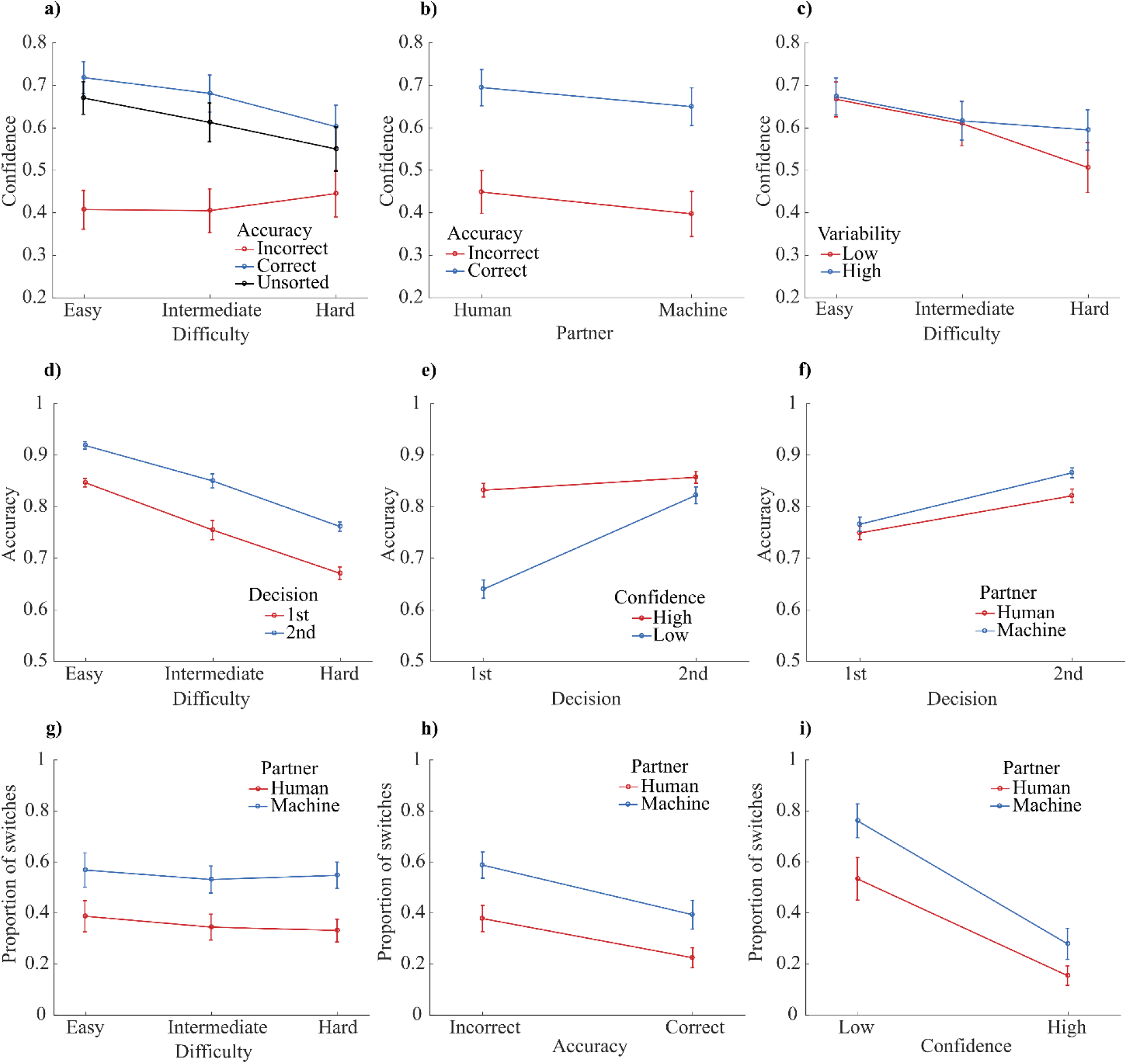
Plots showing dependent variables averaged across participants, with error bars indicating the standard error of the means. (a-c) Proportion of high confidence trials is shown as a function of accuracy and difficulty (a); as a function of accuracy and partner (b); and as a function of variability and difficulty (c). (d-f) Proportion of correct responses is shown as a function of task difficulty and decision (first and second perceptual decision; d); as a function of confidence and decision (e); and as a function of decision and partner (f). (g-i) Proportion of switches (in conflict trials) is shown as a function of partner and task difficulty (g) as a function of partner and accuracy (h); and as a function of partner and confidence (i).

Interestingly, a main effect of Partner was also found (χ^2^(1) = 6.70, p = 0.01), with lower confidence in the blocks in which participants interacted with a machine (M = 0.59, SD = 0.17) compared to when they interacted with a human partner (M = 0.63, SD = 0.17), and irrespectively of whether the response was correct or incorrect (see Figure 2b).

In sum, confidence strongly correlated with accuracy, showing that participants could clearly estimate the correctness of their perceptual decisions. Confidence was also affected by the identity of the interacting partner, while the overall accuracy (and reaction times; see supplementary materials) remained unaffected by the partner’s identity. Contrary to previous study (Desender et al. 2018) stimulus variability only mildly influenced confidence judgments, that is, only when task difficulty was hard.

### Accuracy

Linear mixed models assessed the impact of partners’ report and confidence on the participants’ accuracy in the two successive perceptual decisions of the task. The model included Decision (1^st^, 2^nd^), Partner (human, machine), Confidence (low, high), Variability (low, high) and Difficulty (hard, intermediate, easy) and the double interactions between Decision and each of these factors^3^. The same factors and interactions were included as uncorrelated by-participants random slopes and intercepts. The analyses showed a main effect of Decision (χ^2^(1) = 51.93, p < 0.001), with higher accuracy in the second decisions, i.e., after viewing the partner’s report (accuracy 1^st^ decision: M = 0.76, SD = 0.04; 2^nd^ decision: M = 0.84, SD = 0.03), and similarly across the different difficulty levels (Figure 2d). The interactions between Decision and Partner (χ^2^(2) = 6.39, p < 0.011), Decision and Confidence (χ^2^(2) = 40.50, p < 0.001), and Decision and Difficulty (χ^2^(2) = 10.71, p = 0.005) were also significant. Post-hoc tests showed that accuracy increased in the second decision both when participants interacted with a human and a machine partner (p < 0.001; see Figure 2f). In addition, as illustrated by Figure 2f, accuracy was higher (p < 0.002) in the second decision when participants interacted with a machine (M = 0.87, SD = 0.035) compared to when they interacted with a human partner (M = 0.82, SD = 0.049), while no effect of Partner was observed for the first perceptual decision. This may suggest that participants tended to switch more often for the partner’s report when they thought they interacted with a machine compared to when they believed to interact with a human (see switch responses below).

Further post-hoc comparisons showed that accuracy was higher in high compared to low confidence trials for the first perceptual decision (p < 0.001; Figure 2e, left side of the plot). No difference in accuracy between low and high confidence trials was observed for the second decision (p = 0.174; Figure 2e, right side of the plot). Moreover, accuracy was higher for the 2^nd^ decision compared to the first only when participants were not confident about their perceptual decision (p < 0.001). This suggests that participants tended to switch for the partners’ response in particular when they were not confident in their decision, which improved their performances (see Figure 2e). Post-hoc comparisons exploring the interaction between Difficulty and Decision showed that accuracy was higher in the second Decision compared to the first for each difficulty level (p < 0.001).

In sum, whether participants decided to keep or change their response depended on the confidence they had on their own perceptual decision and on the identity of the partner. This point is also corroborated by the analyses of switch responses below.

### Switch responses

A linear mixed model assessed the impact of confidence, accuracy and partner on participants’ decisions to keep their initial perceptual response or change it for the partner’s report. The model included Confidence, Accuracy and Partner and their interactions, as well as Variability and difficulty as predictors. The same factors and interactions were used as by-participants uncorrelated random slopes and intercepts. This analysis was performed only on conflict trials, i.e., trials where the participant selected a different response than the one chosen by the partner (in a total of 8064 trials, only once a participant decided to change his/her response even though there was no conflict with the partner’s response). The analysis showed three main effects: a main effect of Partner (χ^2^(2) = 18.80, p < 0.001), with a larger proportion of switches when participants interacted with a machine (M = 54%, SD = 19%) compared to when they interacted with a human partner (M = 35%, SD = 17%); a main effect of Accuracy (χ^2^(1) = 4.31, p = 0.04), showing that participants switched more often for the partner’s report when they were incorrect (Figure 2h); and a large main effect of Confidence (χ^2^(1) = 61.21, p < 0.001), where participants switched more often when they were little confident regarding their response (Figure 2i).

Interestingly, a Wilcoxon signed rank showed that participants tended to change their initial decision more often based on their confidence than their accuracy. Specifically, participants changed their response on average 49.2% more often when they were low confident compared to high confidence trials, while they changed their initial response 16.4% more often when they were incorrect compared to correct trials. These changes in switch responses were similar when considering the trials in which participants interacted with a machine and the trials where they interacted with a human partner, separately. Finally, no effect of difficulty on the proportion of switches was observed (Figure 2g).

### Pupillary response

A cluster-based permutation test showed that pupil velocity observed in high and low confidence trials differed within a time window going from 550 to 1150 ms (Figure 3a). The pupil velocity observed within this time window was averaged and analyzed using linear mixed models. Given the large number of factors and interactions that could show a relationship with pupil dilation, a model comparison approach was used to remove from the model the factors that do not improve model prediction. We started with a complex model including Partner, Variability, Difficulty, Accuracy, Confidence, and their interactions as fixed effects. The model also included Switch response as predictor and the factor Participant as a random intercept. We then compared this model with a simpler model from which we removed all interaction terms. The comparison showed that including the interactions to the model did not significantly improve model fitting χ^2^(41) = 52.39, p = 0.109. Consequently, we continued our analyses with the more parsimonious model (i.e., the model including Partner, Variability, Difficulty, Accuracy, Confidence, Switch responses as fixed effects and Participant as a random intercept). We performed a backward elimination of the remaining fixed factors in order to find the subset of parameters leading to the best performing model. This led to a final model with only Accuracy and Confidence as fixed factors. Specifically, pupil velocity was higher for correct compared to incorrect responses (χ^2^(1) = 17.54, p < 0.001), and this effect was almost 3 times stronger when comparing high and low confidence (χ^2^(1) = 41.15, p < 0.001), with higher dilation velocity for high compared to low confidence judgments.

**Figure 3.**
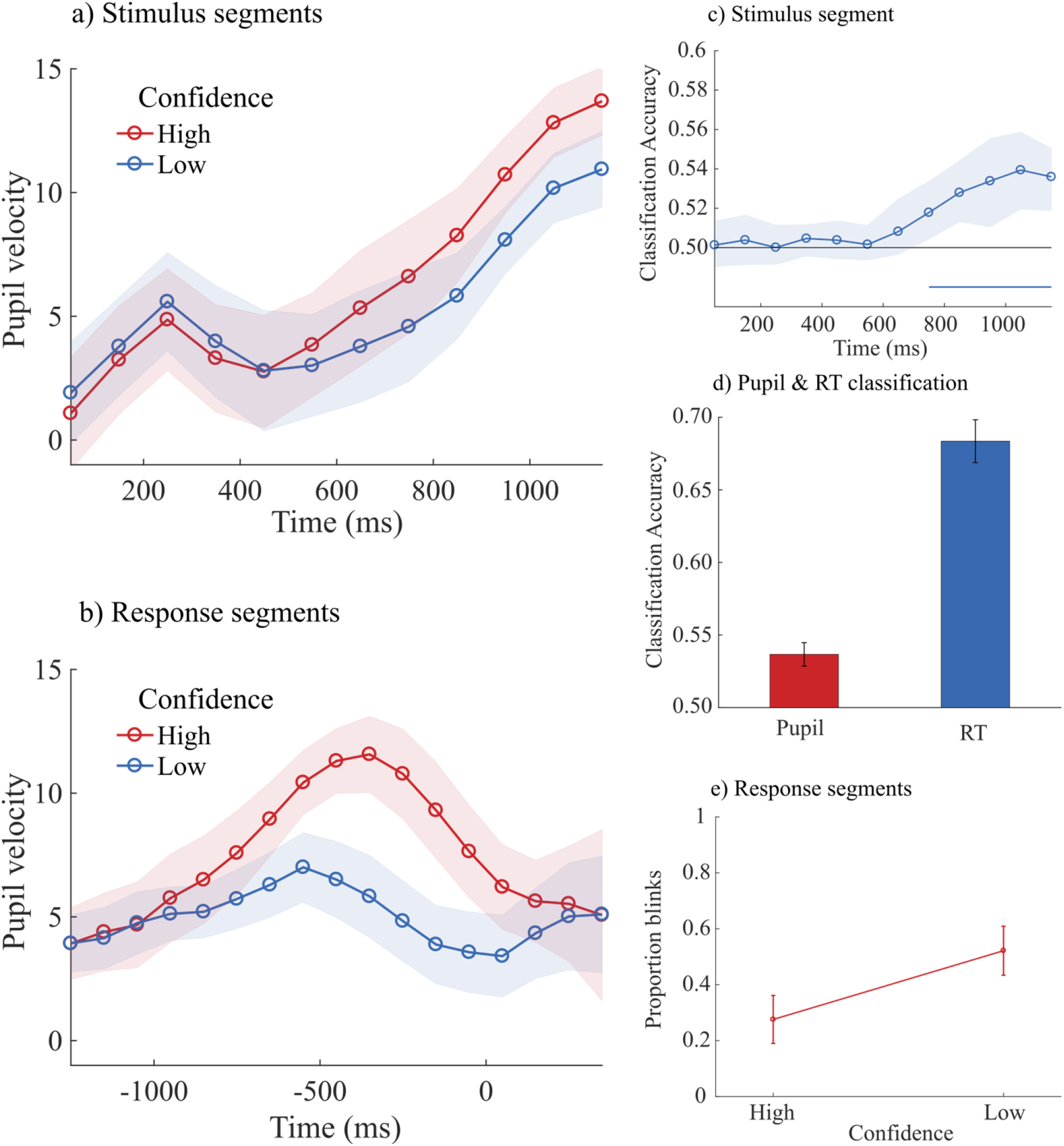
The plots 3a and 3b show the average pupil velocity for high and low confidence trials calculated across participants as a function of the time after stimulus onset (a), and the time around the perceptual decision (b). Shaded areas represent ± 1 standard error of the mean. (c) Accuracy (proportion of correct classification) of a classifier dissociating high and low confidence trials from pupil velocity. The classifier was trained and tested on stimulus-locked segments. The horizontal blue line indicates the time points with a classification accuracy that was significantly above chance level (50%). Shaded area represents bootstrapped 95% confidence intervals. (d) Best classification accuracy (across time) to discriminate high and low confidence from pupil velocity (in red) and from reaction times (in blue). Error bars represent standard errors. (e) Proportion of blinks that occurred in high and low confidence trials.

The same analyses on response-locked segments showed a difference between high and low confidence trials within a time window going from -550 to 50 ms (Figure 3b). Pupil velocity observed within this time window was averaged and analyzed using the same approach described above. Including all the interaction terms in the model did not improve model fitting χ^2^(41) = 41.75, p = 0.438. Analyses were then continued with a parsimonious model (i.e., the model including Partner, Variability, Difficulty, Accuracy, Confidence, Switch responses as fixed effects and Participant as a random intercept). The best performing model included Partner, Difficulty, Confidence, Accuracy and Switch response as fixed predictors. Specifically, pupil dilation velocity was higher when participants interacted with a machine compared to when they interacted with a human partner (χ^2^(1) = 10.08, p = 0.002), when their response was correct compared to incorrect responses (χ^2^(1) = 18.94, p < 0.001), when they did not change their initial response (χ^2^(1) = 5.83, p = 0.016), when the task was easy compared to hard trials (χ^2^(2) = 6.41, p = 0.041), and finally more than the other factors, pupil dilation velocity predicted very strongly individuals’ confidence judgments (χ^2^(1) = 154.77, p < 0.001), with faster pupil dilation for high compared to low confidence judgments.

### Classification of pupil and RT

Similarly, to the analyses reported above, pupil-based classifiers could predict significantly above chance level individuals’ judgments of confidence within a time window going from 750 to 1150 ms after stimulus onset (Figure 3c, see also supplementary material for additional classification analyses). The best performance (across time) of the pupil classifier was 54%. RT classifier could also predict significantly above chance level individual’s judgments of confidence with an average accuracy of 69% (Figure 3d). A Wilcoxon Rank Sum Test showed that classification accuracy was higher with RT compared to the classification accuracy observed with pupil velocity z(13) = 4.434, p < 0.001.

### Blink data

Changes in pupil dilation may partly be caused by eye blinks (Yoo et al., 2021). The response-locked segments contained eye blinks, hence we decided to verify whether the proportion of blinks differed across conditions. The number of blinks observed in the response-locked epochs were analyzed with a mixed linear model including Partner, Difficulty, Confidence, Accuracy and Switch response as fixed predictors (i.e., the same final model used for pupil analyses in the same segments), participants was included as a random intercept. The number of blinks increased with incorrect compared to correct responses χ^2^(1) = 7.45, p = 0.006, it also increased when participants changed their initial response χ^2^(1) = 14.80, p < 0.001, and when their confidence was low compared to high confidence trials, χ^2^(1) = 308.14, p < 0.001 (Figure 3e).

## Discussion

This study aimed at contributing to i) the understanding of the impact of confidence in post-decisional behavior, ii) explore the link between pre-response pupil dilation and confidence, and iii) confront personal perceptual decisions to the ones of other humans and machines. Participants completed a 2-alternative-forced-choice discrimination task where they had to identify the dot display containing a dot motion direction that was closer to the vertical axis. After their perceptual decision they estimated how confident they were. They then viewed in different blocks the response of a human or a machine partner. They were told to evaluate the response of their partner and use it to improve their own accuracy on the perceptual task. Specifically, participants could either keep their initial perceptual decision or change it for the other option. Concomitantly, we recorded participants’ pupil dilation to investigate whether it was a good predictor of confidence and whether it was informative about individuals’ strategies to keep or change their initial perceptual response.

In agreement with past research on visual confidence, we observed that participants could evaluate the correctness of their visual decision and that confidence was sensitive to the quality of sensory information (cf. Mamassian 2016 for a review). In addition, we found that interacting with a machine or a human partner modulated confidence judgments without impacting decision accuracy. In particular, when participants interacted with a machine, they exhibited lower confidence judgments in their initial decision compared to when they interacted with a human partner. In line with recent models of metacognition (Fleming & Lau, 2014; Mamassian, 2016), this suggests that confidence judgments do not rely only on the quality of current sensory evidence, but also on additional pieces of information that can bias confidence evaluations (Shekhar & Rahnev, 2020) such as prior beliefs or contextual information (Fleming et al., 2018; Desender et al., 2022). Interestingly, the decrease in confidence that participants exhibited when interacting with a machine was accompanied with a larger tendency to change their initial perceptual responses, thus boosting participants’ final performance accuracy. This seems to suggest that independently of accuracy and task difficulty, confidence estimations drove participants’ strategies to keep or change their initial perceptual judgments. A possibility is that, even though the performance accuracy of the two fictive partners was strictly the same across the task, participants may have had prior assumptions that the machine would be more reliable than a human on the task. Hence, to achieve better performances from their subjective perspective, participants may attribute less weight in their own responses compared to those given by the machine through a decrease in the confidence associated with their own responses, which may in turn enable them to adapt their post-decisional strategies by modulating the likelihood of a decision switch (Mattingly et al., 2016). The evidence for a modulation of confidence based on the identity of the partner is interesting. However, it is important to underline that the direction of this effect should be taken with caution, as it may originate from personal considerations and contextual factors. In line with this, recent studies showed that machines inspire overconfidence (Booth, 2017; 2020) or mistrust (Nicodeme, 2020; Lee & Rich, 2021; Seth et al., 2020) depending on the situation. Furthermore, it has been shown that prior beliefs about a task could induce under- and overconfidence (Van Marcke et al., 2022), and that confidence plays a role in shaping certain aspects of decision-making behavior such as the confirmation bias (Rollwage et al., 2020) as well as driving post-decisional behaviors such as changes of mind (Rollwage et al., 2020; Pescetelli et al., 2021) or decision switch (Mattingly et al., 2016). An alternative hypothesis is that the belief that the machine would perform better would lead participants to pay less attention to the perceptual task and strongly rely on the machine’s answers. However, this seems unlikely since no difference in accuracy in their initial perceptual decision was observed when participants interacted with a machine compared to when they interacted with a human partner.

Further evidence strengthens the interpretation of a key role for confidence in guiding post-decisional behavior. Specifically, participants changed their initial response approximately 49% of the time when they were not confident compared to when they were confident, while around 16% of the time when their response was incorrect compared to when it was correct. Importantly, this behavior was observed both when they interacted with a machine or a human partner. Taken together these findings suggest that subjective confidence estimates strongly impact post-decisional behavior, more than perceptual accuracy or task difficulty. Indeed, we observed no modulation of task difficulty on the decision to keep or change initial responses, reinforcing the idea that what matters for subsequent behavior is the subjective representation of decision uncertainty (i.e., confidence). This seems obvious considering the fact that the brain cannot have direct access to objective external information, and must also take into account other sources of information such as internal states and prior knowledge. Confidence may well be one of the cognitive products of multi-dimensional integrated information used to update their decisions.

Our study provides evidence supporting that confidence plays a role in decisional behaviors, thus corroborating recent studies suggesting that confidence guides information seeking and learning (Desender et al., 2018; Meyniel et al., 2015; Guggenmos et al., 2016) as well as change of mind (Fleming et al., 2018, Rollwage et al., 2020; Pescetelli et al., 2021) and decision switch (Mattingly et al., 2016), and it may be one of the mechanisms involved in human-human (Bang et al., 2017; De Martino et al., 2017) and human-machine interactions (Wright et al., 2020; Zhang et al., 2020).

Contrary to previous studies, confidence was weakly modulated by the variability of the stimulus (Desender et al., 2018). This could be explained by interindividual differences. Indeed, some participants may prefer stimuli with high variability while others stimuli with low variability stimuli (cf. de Gardelle & Mamassian, 2015).

Decision-making is accompanied by broad neurophysiological changes of the body (O’Connell & Kelly, 2021) including changes in eye pupil activity (Urai et al. 2017). In the present study we also investigated the relation between eye pupil changes and confidence. In particular, we analyzed pupil velocity since recent studies suggest that temporal derivative of pupil dilation reflects closely arousal fluctuations (Crombie et al., 2021; Okun et al., 2019; Reimer et al., 2014, 2016). The interest in using pupil dilation as a proxy of confidence relies on the fact that it could allow for the monitoring of individuals’ confidence and uncertainty online and through time without the need of collecting confidence judgments or measuring response times. This application can be important in different domains, including the field of human-human and human-machine interactions. There is indeed growing interest in neuroergonomics to monitor through time different cognitive states of operators while they interact with technology and their teammates (Dehais et al., 2017, 2020; Gramann et al., 2017), with the objective among others to conceive and evaluate new technology. In this context confidence appears to be a critical phenomenon for optimal human-machine interactions (Lee & Moray, 1992; Parasuraman et al., 1993), and to that extent, online access to operators’ confidence states could be particularly useful.

Interestingly, in agreement with previous studies (e.g., Lempert et al., 2015) confidence correlated with changes in pupil dilation dynamics. However, in addition to past research, our study showed that pre-response pupil dynamics predict confidence judgments. Specifically, we observed that during the presentation of the visual stimulus (when neither blinks nor responses could occur) pupil velocity was higher when the following response was correct or confident. Interestingly, pupil velocity did not correlate with the difficulty of the task. This dissociation together with the strong relation between pupil velocity and confidence (three times stronger than the relation between accuracy and pupil velocity) suggest that pupil velocity may be associated with the subjective evaluation of uncertainty rather than objective uncertainty, through short-term arousal fluctuations. A possible explanation for this finding would be that in confident trials participants allocated more strongly attentional resources to the stimulus compared to low confidence trials. However, if that was the case, we believe that pupil dynamics would then correlate more with correct responses than confidence. Our finding showed exactly the opposite: larger variations of pupil dynamics were observed when confidence varied rather than when accuracy varied. Secondly, pupil dilation dynamics differed between high and low confident trials also when focusing only on correct trials (see supplementary materials). In agreement with confidence-based learning models, another possible explanation is that confidence has a mechanistic role in reward and value-based learning (Guggenmos et al., 2016; Ptasczynski et al., 2022). Interestingly, pupil dilatation, and more generally arousal, is associated with reward anticipation (Schneider et al., 2018; Koelewijn et al., 2018). We speculate that feeling confident of one’s own performance during a task may work as a reinforcing and rewarding signal influencing cognitive states, which would in turn modulate arousal-based pupil fluctuations.

Past research reported relations between pupil dilation and uncertainty post-response (Lempert et al., 2015; Urai et al., 2017; Balsdon et al., 2020). Our study provides new evidence on the link between pre-response pupil changes and confidence. In further analyses we found that pupil dilations predict confidence only weakly (54% correct classification) compared to response latencies related to perceptual decisions (69% correct classification). Hence, reaction time remains a rather reliable proxy of confidence estimations, in line with previous literature (Kiani, 2014). However, in the absence of any overt behavior, pupil dilations may provide useful insight in the confidence state of an individual, better than a random classification that is at 50%. Further investigation should be encouraged to better understand the relationship between pupil and confidence and to improve classification accuracy of pupil-based confidence signals.

The analyses of pupil changes around the time of the perceptual response, provided similar findings showing that, starting from -550 ms to +50 ms around the response, pupil velocity correlated strongly with confidence, accuracy, task difficulty and participants’ tendency to change their initial response. However, this time window was contaminated by eye blinks which may have driven unwanted changes in pupil dilation velocity, thus making any conclusion regarding the changes in pupil velocity observed during this time period difficult to draw. In fact, we observed a larger proportion of blinks in hard, incorrect and in low confidence trials. It is well known that stress, boredom, and fatigue can induce an increased blink rate (Tanaka et al., 1993; Barbato et al., 2000; Danckert et al., 2018). We speculate that the increase in eye blinks observed either in low confidence trials or in incorrect and hard trials are associated with momentary stress changes associated with low performance and task difficulty. Further studies should corroborate the relationship between confidence and eye blinks.

## Conclusions

In summary, our findings bring new evidence supporting that confidence contributes to post-decisional behaviors and in particular decision switches during the interaction with another partner. Confidence would be a subjective evaluation of decision uncertainty, which integrates internal representations including current internal states and context-dependent beliefs about ourselves and others (Lebreton et al., 2019; Carlebach et al., 2023) in situations with incomplete knowledge (Khalvati et al., 2021). The role of confidence may be to weigh our decisions and perceptions, allowing adaptation and learning, therefore contributing to decisional behaviors. Our results also provide new insight on the relation between eye pupil dilation and confidence. Crucially, eye pupil dynamics seem to provide online information about ongoing metacognitive processes and could also directly participate in decision-making and metacognition, in line with recent advances in the neurophysiology of perceptual decision-making (O’Connell & Kelly, 2021).

## Acknowledgment

This research was funded by the Agence Nationale de la Recherche, research funding ANR JCJC, project number ANR-18-CE10-0001, awarded to Andrea Desantis.

Data set will be made available on osf.io/h64xs.

## Footnotes

1 The filter amplitude was determined as 1 divided by twice the average trial duration. On average a trial lasted 6 sec (including the 0.5 sec of ITI). This was done to remove slow oscillations that were not linked to our manipulations

2 With the EyeLink 1000 Plus, pupil area is calculated as the sum of the number of pixels inside the detected pupil contour.

3 A full model including the full interaction was also tested, but this model did not significantly improve the model fitting compared to the simpler model.

## Notes

### Competing Interest Statement

The authors have declared no competing interest.

### Summary of Updates

Some minor updates on the text and figures after review of our first journal submission.

