## Supplementary Material for "What confidence and the eyes can tell about interacting with a partner"

### The role of confidence in decision-making and its pupillary correlate during the interaction with a partner

**Results**

*Reaction Times.* The same analyses on RTs showed a main effect of Difficulty ($\chi2(2) = 19.18$, $p < 0.001$), with gradually faster perceptual responses with the decrease of task difficulty (hard trials: M = 1.896, SD = 0.189; intermediate trials: M = 1.850, SD = 0.187; easy trials: M = 1.790, SD = 0.172). In addition, we observed a significant interaction between Partner and Difficulty ($\chi2(2) = 6.94$, $p = 0.031$) and between Variance and Difficulty ($\chi2(2) = 10.01$, $p < 0.007$). None of the other comparisons were significant. Simple main effects analyses investigated the two significant interactions. In particular, we compared human and machine partner trials for each difficulty level. The analyses showed that RTs were faster only in hard trials ($p = 0.025$) when participants interacted with a human partner (M = 1.847, SD = 0.153) compared to when they interacted with a machine (M = 1.945, SD = 0.264). In other words, decisions in hard trials may have been taken faster when people interacted with a human compared to a machine partner. Finally, we compared low and high variance trials for each difficulty level. RTs were faster in hard trials ($p = 0.003$) with high variance (M = 1.852, SD = 0.166) compared hard trials with low variance (M = 1.940 SD = 0.220). Accordingly, decisions may have been taken faster when participants performed hard trials with high variance than with low variance.

*Classification results.* The blue curve in Figure 1 depicts the accuracy (proportion of correct classification) of a classifier trained to differentiate, from pupil dilation, high and low confidence judgments in correct responses only. The classifier was trained and tested on stimulus-locked segments. The horizontal blue line depicts the time cluster with classification accuracy significantly above chance level (50%). Shaded area represents bootstrapped 95% confidence intervals. The red curve depicts the accuracy of a classifier trained to dissociate correct and incorrect responses in high confident trials only from pupil dilation. No significant time cluster was observed for the latter classifier.

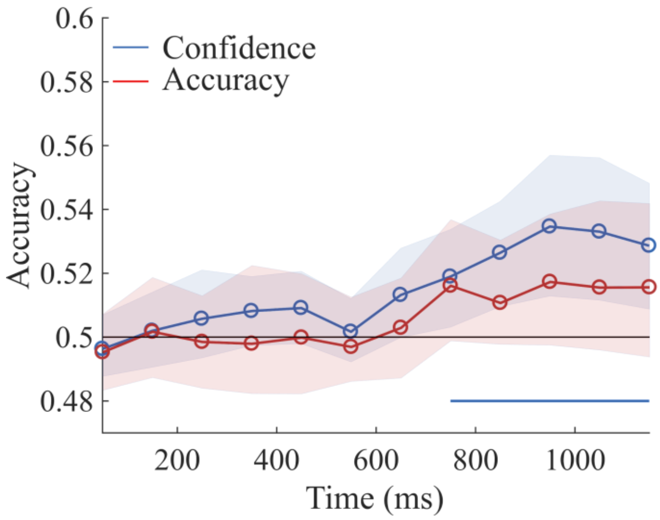

Figure 1

At the end of the experiment, each participant was asked to answer a series of questions (Table 1) addressing his/her interaction with the computer and human partner. The main objective of this questionnaire was to evaluate whether participants questioned the fact that they were really viewing the responses or another participant or a machine-learning program. From the questionnaire and a short debriefing, participants trusted the scenario we created for these experiments. About 56% of participants felt that the computer partner performed better than the human partner and they felt they used the former's responses more often. About 31% of participants said they could not see or be able to differentiate the performance of the two types of partners.

Table1. Questionnaire participants filled at the end of the experiment. English translation from French.

| **Questions** | **Yes** | **No** | **I don't know** |
|---|---|---|---|
| Did you use the human partner's responses to change your choices? | | | |
| Did you use the computer partner's responses to change your choices? | | | |
| Do you think one partner was better than the other? | | | |
| Do you think that the computer partner was better than the human partner? | | | |
| Did you use more often the responses of one partner than the other to change your choices? | | | |
| Did you always have in mind what type of participant you were interacting with, human or computer? | | | |
